# Real-time monitoring of bacterial growth and fast antimicrobial susceptibility tests exploiting multiple light scattering

**DOI:** 10.1101/481184

**Authors:** SeungYun Han, HyunJung Kim, Jongchan Park, SangYun Lee, KyeoReh Lee, Ju-Kang Kim, Hyun Jung Chung, YongKeun Park

## Abstract

Antimicrobial susceptibility test (AST) is widely used to provide the minimum inhibitory concentration of bacteria, and crucial to provide appropriate uses of antibiotics and to address the issue of drug-resistance bacteria. However, ASTs require the time-consuming incubation about 16-20 h for the visual determination of the growth of bacterial colonies, which has been a major obstacle to on-site applications of ASTs. In this study, we propose a rapid and non-invasive method based on laser speckles to evaluate the bacterial growth movements in real time, thus reducing the time for the agar dilution method. With a simple configuration compatible with conventional agar plates, the analysis of laser speckle from samples enables the early detection of the presence of growth as well as its detailed history of the colony-forming movement on agar plates. Using the samples prepared through the same procedure as the agar dilution method, we obtained the AST results at least 4-8 hours earlier than the conventional method without compromising the accuracy. This technique does not require for the use of exogenous agents, but works for most bacteria regardless of their species. Furthermore, the distinctive responses of several species to microbial agents were revealed through the present technique supporting a comprehensive analysis of the effect of the antibiotics. The findings suggest that this new method could be a useful tool for rapid, simple, and low-cost ASTs in addition to providing the historical information of the bacterial growth on agar plates.

## Introduction

Along with the popular uses of antibiotics for the disease treatment, its misuse and overuse have caused the increase of microbial resistance, threatening the efficacy of antibiotics to bacteria (1). The primary solution is to select appropriate drugs and proper doses to pathogens, which can be achieved through antimicrobial susceptibility tests (ASTs). The gold standards for ASTs are microdilution and agar dilution method (2, 3). These methods observe the bacterial growth at given antibiotic concentrations, in liquid and on solid, respectively. The microdilution method evaluates the bacterial growth measuring the optical density (OD), while the agar dilution method evaluates the visible growth of bacterial colonies with human eyes. Although they offer reliable results with simple procedures, however, they require a long lead-time of 16-20 h (2). This long process is mainly due to the time for bacteria to grow into a substantial size or density to allow its detection. For a microdilution method, the OD measurement requires a minimum bacterial concentration of 10^7^ CFU/ml (4). This high threshold for OD measurement sets limits to not only the microdilution method but also ASTs via automated microdilution systems, such as Vitek2, MicroScan, and BD Pheonix™ (3, 5-9), because it requires the time for test samples to grow into the high concentrations. Likewise, an agar dilution method needs bacterial colonies to develop into a visible size, which requires overnight incubation. The time-consuming procedure for ASTs has been the hurdle to utilize ASTs as guidance for treatments in an early stage. Thus physicians have chosen the initial prescription based on their empirical guess. In need for a faster way to achieve susceptibility test results, other fast diagnostic techniques for ASTs has been developed for decades. Various types of approaches have been introduced, such as flow cytometry (10-12), microfluidic devices (13-17), biofluorescence (18-20), nanoparticles (21), cell growth imaging (22, 23), and magnetic bead rotation (24). However, although they provide powerful methods for fast ASTs, their special instruments, high cost, or need for well-trained technicians often hinder their widespread use. Still, the microdilution and agar dilution methods are the most common tools for ASTs due to its simplicity, low price, and reliability, even though they require a relatively long lead-time. We therefore assume that a faster diagnosis sharing the same procedures of the standard AST methods would give a more substantial impact.

Here, we present a new concept of a speckle-based optical method for bacterial colony growth detection, which can reduce the time required for the agar dilution methods. We utilized the property of laser scattering in highly turbid media. When a temporally coherent laser beam passes through a medium with highly inhomogeneous distributions of refractive indexes (RIs), such as bacteria, the scattered light forms a highly complex intensity distribution, called a speckle pattern, resulted from the interference of multiple light scattering. Because small changes in the RI distribution cause significant alterations in detectable intensity patterns, the sensitivity of a laser speckle patterns scattered from transparent bacteria provides a non-invasive way to inspect the microscopic dynamics of the bacteria on agar plates. Thus, the colony forming movement can be effectively detected in real time. Previously, the speckle field imaging has been used for blood flow visualization (25-28), label-free bacterial colony phenotyping (29), and food condition evaluation (30, 31). Speckle correlation techniques have also been used for several types of sensors (32, 33), which require exceptional sensitivity. Although speckle illumination enhances the contrast for detecting transparent microscopic objects (34, 35), the use of multiple light scattering can greatly enhance the detection sensitivity (36), because the number of interaction between light and target objects dramatically increases. More recently, the use of multiple light scattering has been exploited for the detection of micro-organisms in food, the sensitivity of spectroscopy (37-39).

In this paper, we present a method for the rapid and highly sensitive detection of bacterial growth exploiting multiple light scattering. An agar plate is placed between two optical diffusers, and multiple light scattering is generated when a coherent laser beam was illuminated. The resultant speckle patterns from the plate was recorded and analyzed using a temporal correlation. This correlation analysis serves as an indicator of the movements of bacteria, which is highly related to bacterial growth. The proposed technique not only confirms the presence of bacterial growth, but also provides the detailed characteristics of bacterial growth, including each species, concentrations, and under various antibiotic conditions. Consequently, we demonstrated simple and rapid ASTs with the proposed method. With the high sensitivity of the speckle intensity correlations under bacterial movements, the bacterial growth was detected earlier than the formation of the visible colonies, at most in 12 hours, depending on species. This previously unreported technique allows us to track the growing movement on bacterial colonies in real time, and accomplish ASTs within a few hours.

## Results

### Principle and optical setup.

The principle and schematic of the setup are shown in Fig. 1a. The proposed device examines temporal changes of the speckle field. The speckle intensity distribution is monitored over time in order to analyze its temporal variation. Then the analyzed results explain the movement history in samples. In the absence of bacterial movement as in the left of Fig. 1a, the obtained speckle movie shows little fluctuation over time because of the deterministic nature of light scatterings. On the other hand, with active bacteria on agar plates as in the right of Fig. 1b, the colony-forming movement induces intensive dynamics in samples. This dynamics, perturbing the light paths of the laser beam, causes a large temporal fluctuation.

**Fig. 1.**
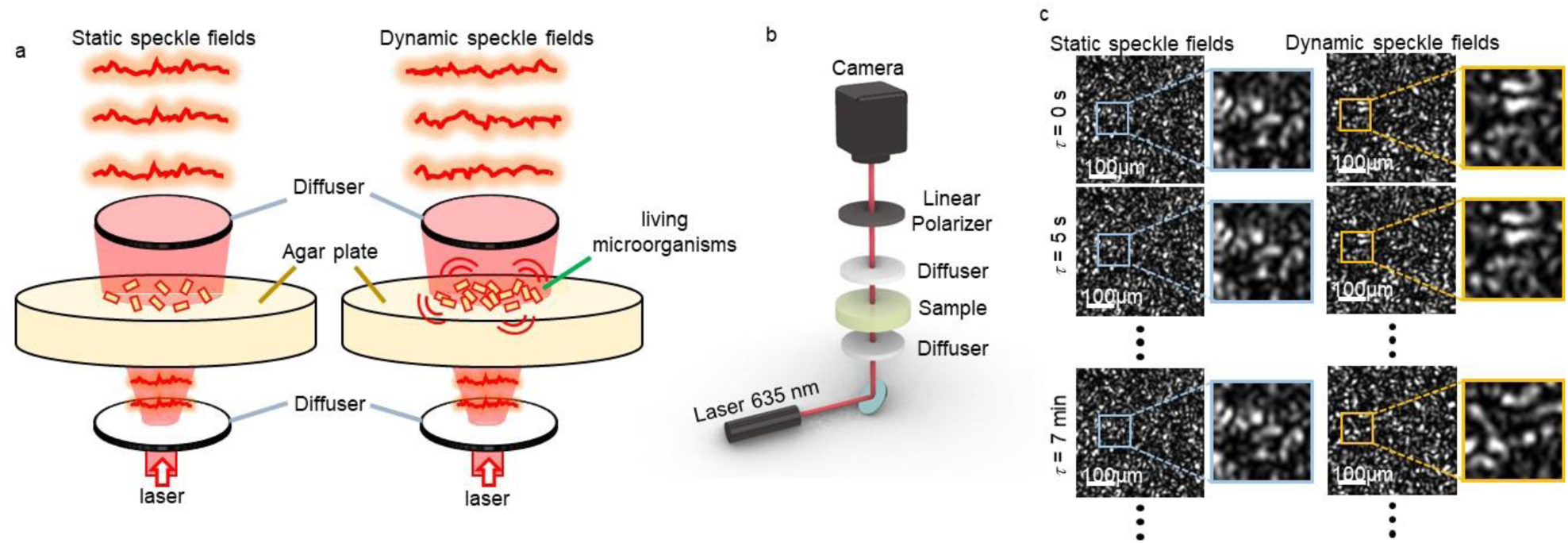
Principle and schematic representation of the proposed optical setup. a, the speckle field is static in the absence of bacterial movement on an agar plate, while dynamic in the presence of the bacterial growth movement. b, the schematic of the optical setup. c, the speckle images in case of static and dynamic samples. The images with static fields have little change over time, while intensively changing with the dynamic sample.

The detailed experimental setup is shown in Fig. 1b. A laser beam from a diode laser module passes a diffuser, an agar plate, and another diffuser in sequence (see Methods for detailed experimental setup). The beam size before the first diffuser is about 1/4 inches and the light precisely illuminated the bacterial area on agar plates. A linear polarizer is added in front of the CCD camera in order to maximize the contrast by the interference. All the experiments are conducted on an optical and the setup is delicately manufactured to minimize any mechanical noise from environments (see Supplementary Fig. 1).

Due to the microscopic size of bacteria, it is difficult to measure the tiny bacterial signal when located inside a bulk volume. To overcome this problem, we used the advantages of exploiting multiple light scattering. Light scatters and interacts with a sample with multiple times due to the presence of two diffusers. Then, the generated speckle field is extremely sensitive to the RI distribution of a sample. Individual bacteria has 3D internal RI distributions which are distinct from that of surrounding media (40, 41). This sensitivity of the speckle field allows the detection of the small bacterial movement by amplifying the signal from the RI change in the sample. This aspect is the main difference from speckle illumination techniques, which only use single light scattering from the samples (34, 35-42).

The speckle images in Fig. 1c are the results from samples made with culture media and 1×10^8^ CFU/ml *E*. *coli* solution, respectively (see Methods section for details). In the absence of bacteria, as expected, the speckle images had little changes over time (see Supplementary Movie 1). In contrast, with the bacterial solution, the speckle images exhibit a significant change after seven minutes (see Supplementary Movie 2). We should note that the change of dynamic speckle pattern occurred in the time scale of several minutes, but not in five seconds. This ten-minute-scale order in temporal change makes coincidence with the growth characteristic of the bacteria, of which cell cycle period is known as twenty minutes (43). Other target bacterial strains used in this paper, *P*. *aeruginosa* and *S*. *aureus*, have similar cell cycle period (44, 45), thus investigating changes during seven minutes is reasonable with the similar timescale.

### Growth movements of bacterial colonies.

Prior to the speckle-based detection, we investigated the colony forming progress via a phase contrast microscope (Fig. 2). An *Escherichia coli* (ATCC 8739) solution = of 1×10^8^ CFU/ml is prepared and 5 μl of this solution, 5×10^5^ CFU in total, is dropped on at the center of an agar plate. By spreading the drop to have a diameter about 2 cm, the CFU per unit area is to set to be similar to the conventional agar dilution protocol (2), which requires 10^4^ CFU for the area of diameter 5 mm. The agar plate is dried in a clean bench for an hour in order to minimize the instability of the wet surface, and then the microscope movies are recorded while the sample is incubated at 35°C with a temperature controller (see Supplementary Movie 3). As illustrated in Fig. 2a, the movement in the sample differs depending on its growing phase. Undergoing a lag phase and an exponential phase, the movement steadily accelerates, and gradually slows down through the end of the exponential phase and the stationary phase. This behavior fully corresponds to the theoretical models (46), which state the growth curve can be described by a sigmoid function. The images in Fig. 2b describe the movement in each phase in a microscopic view, and Fig. 2c shows the pictures of the sample at the corresponding times. The surface of the agar plate is mostly covered by the bacterial colonies in the exponential phase. However, with bare eyes, the colonies can be seen only after reaching the stationary phase.

**Fig. 2.**
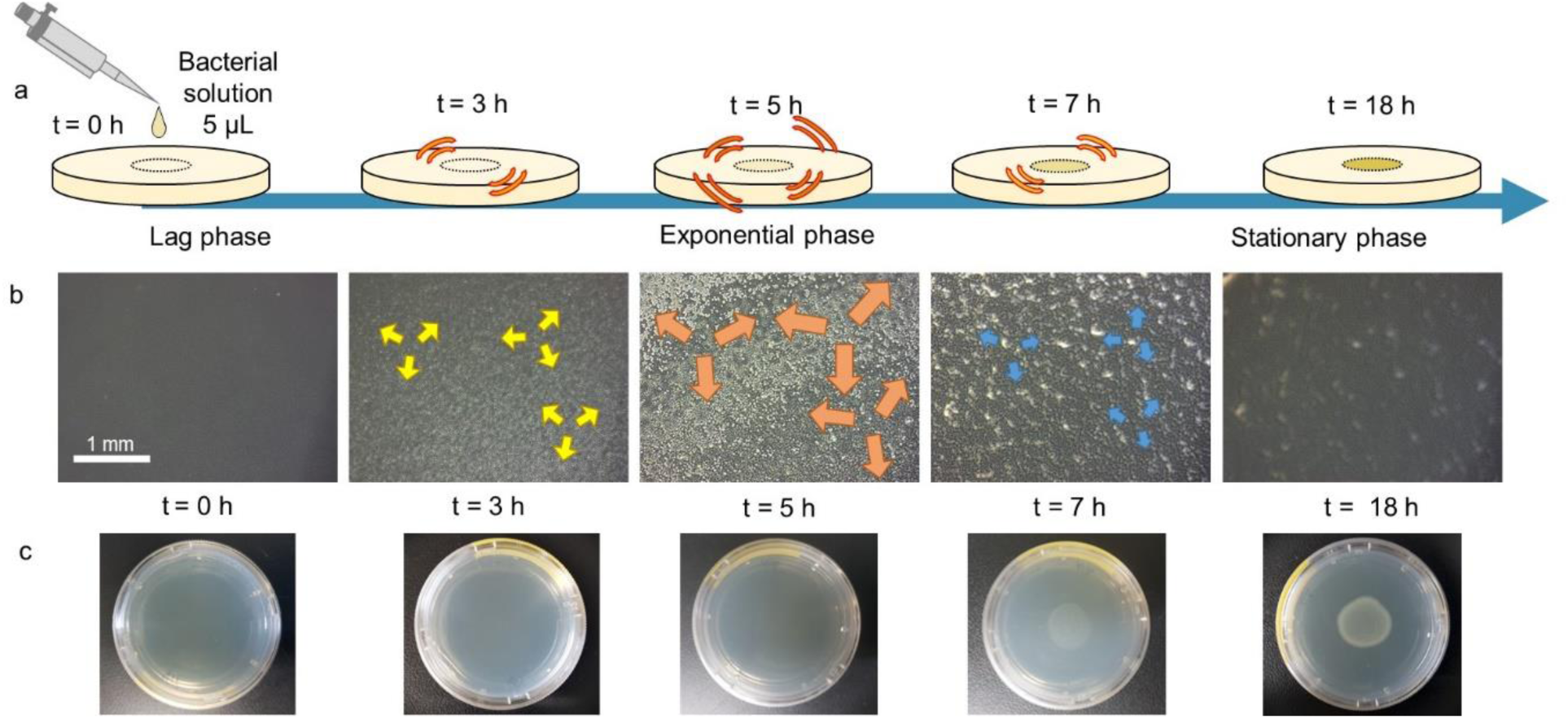
During the development of bacterial colonies, the amount of movement depends on the growth phase. a, a droplet of bacterial solution (*E*. *coli*, ATCC 8739, 1×10^8^ CFU/ml) on an agar plate grows into colonies. The growth movement accelerates from the lag phase to the exponential phase (5 h), then slows down and stops reaching the stationary phase (18 h). b, the microscopic pictures of the bacterial samples. The bacterial area expands and fully covers the surface around the exponential phase, and forms denser colonies until the stationary phase. The arrows in the pictures indicate that the amount of dynamics at a specific time depends on its phase being most vigorous at *t* = 5 h. c, pictures of the agar plate with bacteria for each time. The colonies are visible only after the end of the exponential phase (7 h).

The growth movement of bacterial colonies in agar plates is examined with the proposed speckle-based method. A sample is identically prepared as in Fig. 2, then the CCD camera recorded consecutive 108 frames for nine minutes (one frame per five seconds) at each time in order to analyze the bacterial movement in the sample. The difference between images during nine minutes represents how much movement occurred at that time. Fig. 3a shows the speckle images at *t* = 3, 5, and 7 h, which are representative times for the lag phase, the exponential phase, and the stationary phase, respectively (Supplementary Movies 4, 5, and 6). As can be seen in the first column, in the lag phase, the images of time interval τ= 7 min had only little intensity difference (*ΔI*) and low standard deviation (*Δσ*) during seven minutes. However, in the exponential phase as in the second column, both the intensity difference and the standard deviation severely increased. In the stationary phase, the third column, again there is only little change as the first column. This implies the bacterial growth thoroughly followed the three phases in order.

**Fig. 3.**
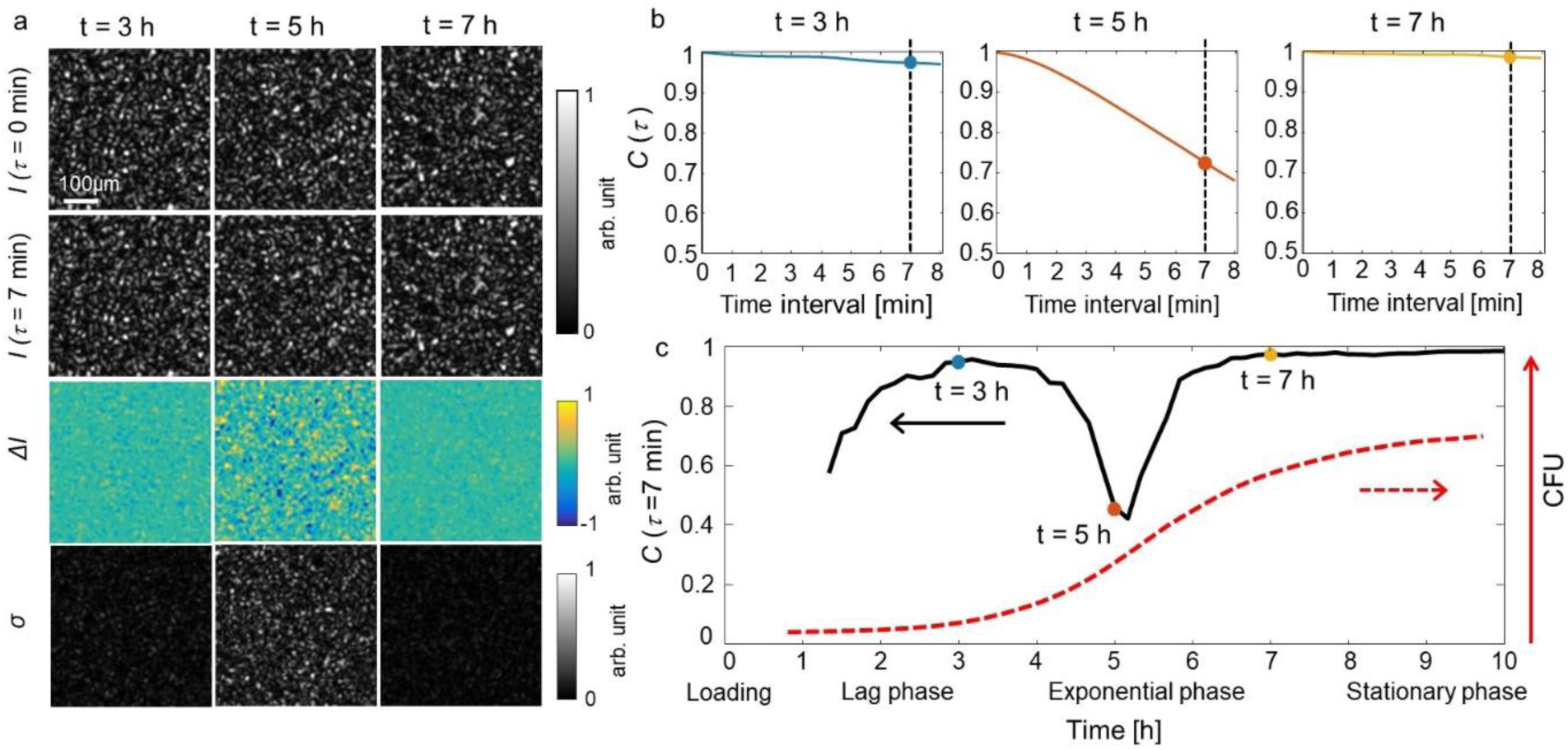
The autocorrelation analysis of the bacterial sample identically prepared as in Fig. 2. a, the dynamics of the speckle movies for each time. For each corresponding time, two images of time interval seven minutes and their intensity difference (*ΔI*), and standard deviation (*σ*) of the movies during seven minutes are presented. Both intensity difference and standard deviation are high at 5 h, which is the exponential phase. b, the autocorrelation graphs of the movies for each time. The graph decreases more rapidly with movies of high temporal changes. c, the history of autocorrelation value at a time interval (*τ*) seven minutes. A reverse peak appears around the exponential phase, indicating the presence of the colonial growth. The total cell number would follow the dotted red line as a sigmoidal function.

### Quantitative analysis of speckle dynamics.

To quantitatively evaluate the growing movement, we calculated the “autocorrelation” for each nine-minute long movies. Before calculation, we need to minimize the shot noise from the camera. Each speckle intensity image was replaced by the average of consecutive ten images starting from the replaced image frame. Although the images during 50 seconds (five seconds for one frame) may have minimal temporal changes, averaging the images is valid given that the time interval we use is longer than one minute (We will take seven minutes for a time interval as discussed below). Then the autocorrelation is calculated by the equation,

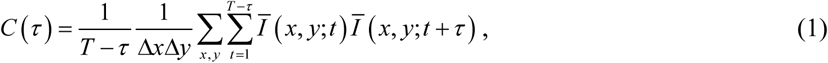

where *Δx*, *Δy* are the numbers of pixels of the CCD camera for *x-*, *y*-direction, *T* is the total time, τis the variable for time interval, and 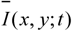 is the normalized intensity,

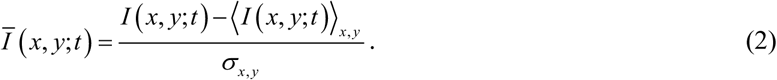

In Eq. (2), ⟨ *I* (*x*, *y*;*t*)⟩ and *σ* _*x*, *y*_ are the average intensity and its standard deviation at position *x*, *y* over time, respectively. The parameter *τ* is from zero to nine minutes in our experiment. The autocorrelation *C*(τ) is the average correlation value between images of time interval τ. Theoretically, the correlation value ranges from 1 to −1 depending on how positively or negatively images are correlated, and is zero for random images. In our experiment, the correlation value remains close to one with samples of no dynamics and zero with samples of vigorous dynamics. Assuming the intensity consists of changing and unchanging portions, *I = I*_*c*_ + *I*_*u*_, the autocorrelation can be divided into two terms,

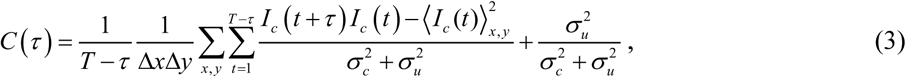

where *σ*_*c*_ and *σ*_*u*_ are standard deviations for changing and unchanging intensity portions, respectively. The first term is the autocorrelation of the changing portion of the intensity, which should be zero by definition. Thus, the *C*(*τ*) value can be written as below.

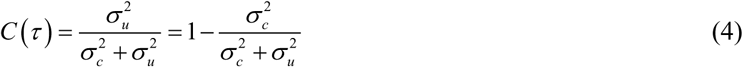

Here we assume that the intensity distribution follows the same statistics regardless of the presence of temporal change; therefore, the decrease in correlation value is in positive relation to the changing portion of light. As the temporally changing portion of intensity is proportional to the area occupied by growing cells, the larger decrease of the correlation value corresponds to the larger amount of movement or, in other words, the bigger number of growing cells. The *C*(τ) graphs at each time are shown in Fig. 3b. The graphs decrease almost linearly as τ increases. This graph shape can be interpreted that the movements in samples occurred constantly without an abrupt shift. However, the slopes at each time are different. The *C*(τ) graphs at *t*_*1*_ = 3 h and *t*_*3*_ = 7 h are almost horizontal, while the graph at *t*_*2*_= 5 h decreases down to 0.7 at *τ* = 7 min from 1 at the beginning, which is relatively steep. This difference is due to the amount of movement in the sample during the lag, exponential, and stationary phases.

For graphical representations, we picked *C*(τ) value at τ = 7 min to represent the movement level in samples at each time. As *C*(τ) linearly decreases with *τ*, the choice of different τ value give the same results, but with different scales without loss of generality. In our demonstration, we simply chose τ = 7 min considering both the number of averaging correlation values and the sufficient decrease of correlation values (for graphs of different τ values, see Supplementary Fig. 2). Fig. 3c shows the *C*(τ=7min) graph during ten hours. This graph was acquired by repeating the speckle movie recording every ten minutes. As shown, the graph starts and ends with the correlation value *C*(τ=7min) close to one, but a reverse peak appears around the exponential phase. To note, the first one hour in *C*(τ=7min) graph is empty because we did not measure during drying (at room temperature) to stabilize the agar surface. The minor increase for two hours at the very beginning is due to the stabilization time after placing the sample on the setup, and this unwanted effect could be minimized using a fresh agar plate and drying it properly. The decrease of correlation value in the exponential phase indicates a vigorous growth movement in the sample. Interpreting the correlation value decrease as the amount of growth, the general shape of the growth curve can be portrayed as the red line of which derivative is the *C*(τ=7min) graph. From the picture of agar plates in Fig. 2c, we can suggest that the colonies become visible between the lowest point in the graph and the end of the peak. It is interesting that the decrease of the correlation value can be identified at the beginning stage of the exponential phase, which is far prior to human eyes can. As the graph can be calculated in real time during the growing progress, the present technique provides an earlier detection for colony formations.

### Correlation graphs from various conditions.

To demonstrate the proposed method can successfully observe colonial growth in real time, proof of principle experiments were performed under various conditions. Speckle intensity patterns from a blank sample and samples of three different species (*Eschericia coli* ATCC 8739, *Pseudomonas aeruginosa* ATCC 9027, and *Staphylococcus aureus* CCARM 3A168) of the concentration of 1×10^8^ CFU/ml were observed. The corresponding *C*(τ=7min) graphs are shown in Fig. 4a. The blank sample, which contains no bacteria, maintains its correlation value close to one throughout the whole experiment time. On the other hand, the three bacterial samples have a decrease in their correlation value, making reverse peaks in their graphs. As mentioned in the previous section, the decreased correlation value is in positive relation to the amount of movement in the sample. The results provide a detailed history and the exact peak times for each species; *E*. *coli* around five hours, *P*. *aeruginosa* around six hours, and *S*. *aureus* around six and eight hours giving two peaks. The presence of two peaks in the correlation graph of *S*. *aureus* was not easily expected because theoretical models for bacterial growth generally expect a smooth sigmoidal growth (46). Previously, these small deviations could have not been reported since all the growth curves for *S*. *aureus* were analyzed based on the typical theoretical models (47, 48). Taking theoretical models as approximations of real situations, the first maximum at *t* = 6 h may be regarded as a small deviation from the larger peak at *t* = 8 h. Nevertheless, it is **inspirational** the proposed method provides precise experimental histories of colonial growth, which may distinguish species by their different graph shapes.

**Fig. 4.**
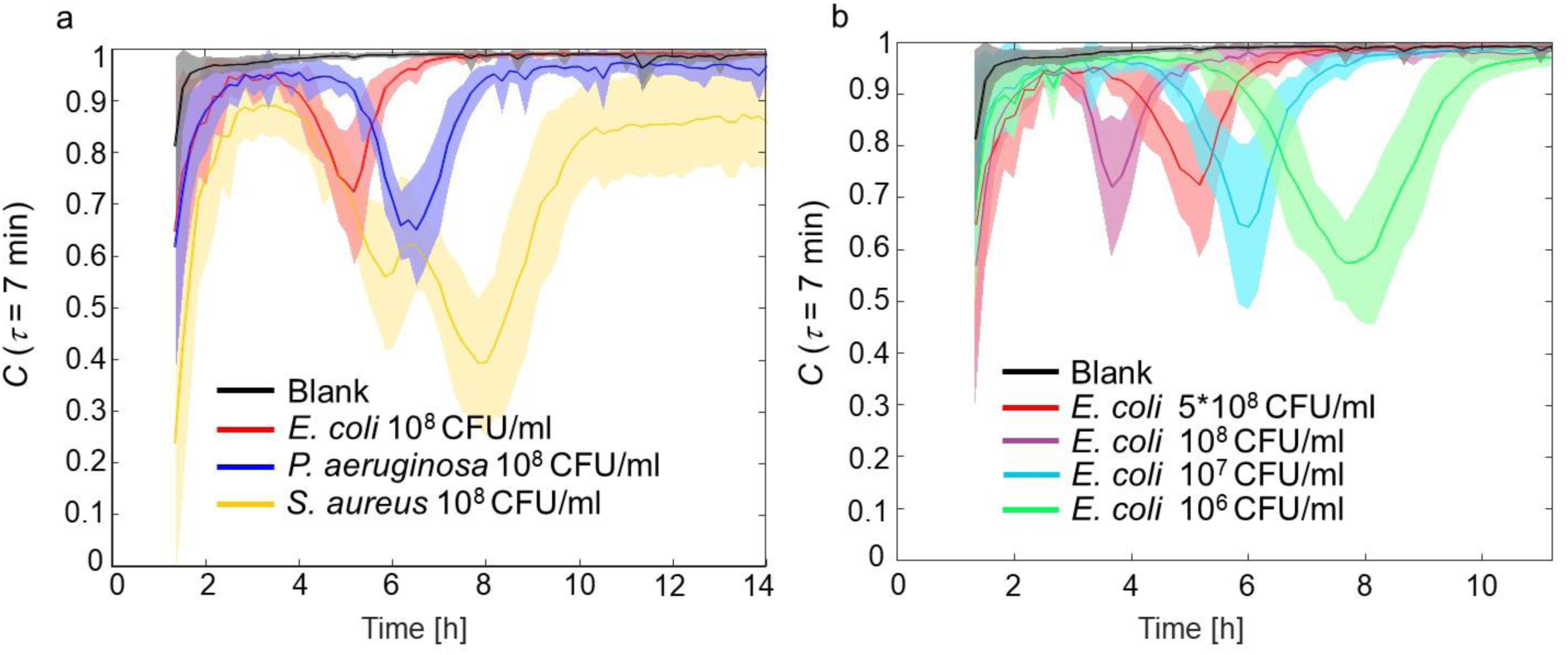
The experimental results with various conditions. a, the C(*τ* = 7 min) graph for different species: *E*. *coli* ATCC 8739, *P*. *aeruginosa* ATCC 9027, and *S*. *aureus* CCARM 3A168 as well as the blank sample (no bacteria). Each line and corresponding shaded area are average and standard deviation, respectively, for ten repeated samples. All samples are from bacterial solutions of concentration 1×10^8^ CFU/ml. The shapes of C(*τ*=7 min) graphs vary depending on the species. b, the C(*τ*=7 min) graph of *E*. *coli* ATCC 8739 with different cell densities. The concentrations of the bacterial solutions used for sample preparations are 5×10^8^, 1×10^8^, 1×10^7^, and 1×10^6^ CFU/ml. The shape of the graph varies depending on the cell density. To note, the reverse peak appears late and become larger as the cell density lessens.

The growth movement was also investigated for different concentrations of *E*. *coli* (ATCC 8739) samples. Samples were prepared as same as in Fig. 2, but with bacterial solutions of different concentrations 5×10^8^, 1×10^8^, 1×10^7^, and 1×10^6^ CFU/ml. The resulting graphs are presented in Fig. 4b. Reverse peaks appear for each of the cases, with the time each peak appears being in the order of concentration. The higher concentration has a peak at an earlier time, while a lower concentration shows a late peak. This difference is mainly due to the period before bacteria fully occupy the surface of agar plates (see Supplementary movie 7 and 8 for different concentrations of growing bacteria). At the beginning of the exponential phase, the sparsely located bacteria grow almost exponentially with sufficient space and nutrient, growth rate being proportional to the cell number. However, as the bacteria grow, the area covered by colonies broadens and eventually nearby colonies collide. The lack of space limits the exponential growth, slowing down the movement in the sample. In this regard, the most vigorous movement occurs just before the collision, and this time depends on the initial surface density of cells because the distance between colonies is different. A lower concentration of bacteria requires a longer time to achieve that certain point, leading to the delay of the peak time. Moreover, the reverse peak gets larger as the concentration becomes lower since the sufficient room for each bacteria accommodates further development. These distinct characteristics for each concentration imply that the cell densities of high-density samples can be determined without dilution if its calibration precedes.

### Rapid AST with current setup.

The proposed technique was utilized to perform ASTs, and the results were compared to the agar dilution method. In these experiments, we used different bacterial strains that have appropriate level of resistance to selected antibiotics. For a robust criterion for AST, the presence of a decreasing trend in correlation value was used as the indicator of bacterial growth. We suggest that the continuous decreasing trend of correlation values, five consecutive points in this work, discriminates the presence of bacterial growth. This criterion is satisfied when there is a clear peak in the correlation graph, *C*(*τ*=7min), and also a moderate decreasing trend in the graph, more than five points, indicated bacterial growth. The MIC values with this criterion well matched to the conventional agar dilution method for three different combinations of bacterial strain and antimicrobial agents. These AST results demonstrated a short time AST was possible with the proposed speckle-based method.

Fig. 5a shows the data from *S*. *aureus* (ATCC 29213) samples on agar plates with *Ampicillin*, where the bacterial concentration of the droplet was 1×10^8^CFU/ml. The minimum inhibitory concentration (MIC) was measured as 2 μg/ml by the agar dilution method, where the agar dilution was performed by investigating the formation of colonies on the sample after one day. In the *C*(τ=7min) graph, the bacterial sample without antibiotics (0 μg/ml) shows a clear peak around *t* = 5 h. The sample of antibiotic concentration 1 μg/ml also has a peak at a similar time *t* = 5 h, although the decrease is relatively small. The peaks indicate the resistance of the bacteria according to the criterion suggested above. However, the samples with antibiotics concentration over MIC, 2 and 4 μg/ml, showed no decrease in correlation value throughout the whole inoculation time, indicating the bacteria’s susceptibility to the antibiotics. This result matched to the MIC result from the agar dilution method, which was also 2 μg/ml.

**Fig. 5.**
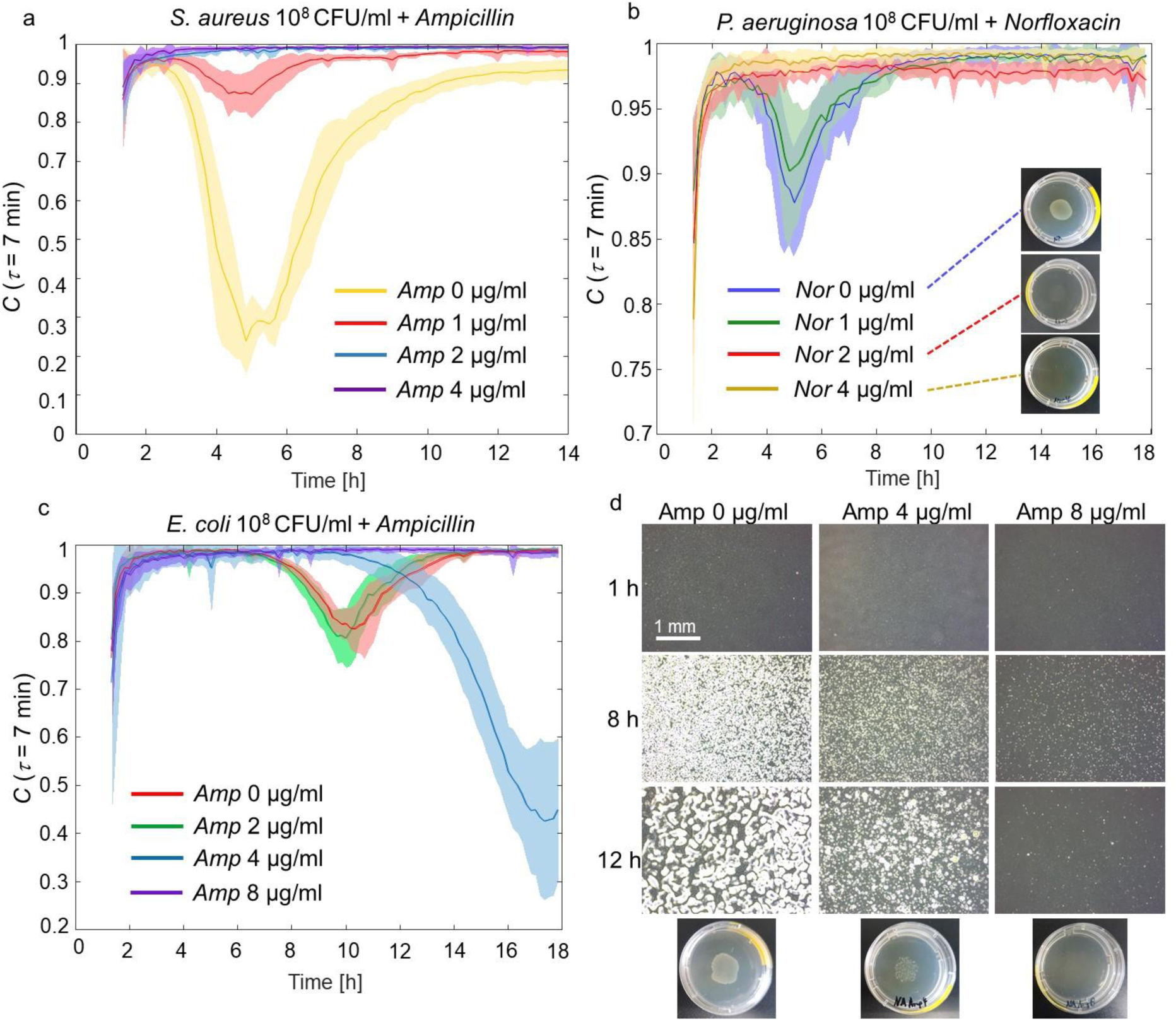
ASTs with the proposed optical setup. Real-time monitoring of the bacterial growth is possible allowing fast ASTs. a, b, c the susceptibility tests of *S*. *aureus* ATCC 29213 to *Ampicillin*, *P*. *aeruginosa* CCARM 0219 to *Norfloxacin*, and *E*. *coli* CCARM 0230 to *Ampicillin*, respectively. Each line is the average and corresponding shaded area is the standard deviation for ten repeated samples. The samples with visible bacterial growth (resistant) showed a decrease in the correlation value. The MIC can be determined as the minimum antibiotic concentration that has no decrease in the correlation graph. The decreasing trend in the correlation value can be identified at least before 12 hours even for slowly growing samples, allowing a shorter test time for the agar dilution method. d microscopic and phone-camera pictures of the samples used in c. The pictures below represent the final colony shapes after one-day incubation. The colonies are sparsely formed in case of *Amp* 4 μg/ml.

The susceptibility test results from other combinations of species and antimicrobial agents showed a different bacterial response to antibiotics. The graph in Fig. 5b shows the *C*(τ=7min) graph for *P*. *aeruginosa* CCARM 0219 to *Norfloxacin*. Up to concentration 1 μg/ml of *Norfloxacin*, a clear peak appears around 5-6 hours, which represent the growth of bacteria, while 4 μg/ml has a nearly constant correlation value close to one indicating no growth. However, the *C*(τ=7min) graph with the concentration 2 μg/ml of *Norfloxacin* has a different shape that a modestly lower level of the correlation value is maintained after 10 hours. According to the criterion suggested above, the decreasing trend over five points is observed around 10 hours, indicating that the bacteria survives and grows in this antibiotic concentration. Therefore, the MIC value with the present method is 4 μg/ml, and this result was consistent with the agar dilution method as presented in the pictures in Fig. 5b. The colony formations were observed up to 2 μg/ml of *Norfloxacin*, yielding the MIC of 4 μg/ml by the agar dilution method. However, the moderate decrease in case of 2 μg/ml can be interpreted as the bacterial response of slow growth in the presence of antibiotics. The suppressed colonies in the picture of 2 μg/ml of *Norfloxacin* supports this result. To note, this strain of *S*. *aureus* here also showed two close peaks around 5-6 hours, similarly as the previous *S*. *aureus* CCARM 3A168 despite of different peak times.

Another AST result of *E*. *coli* (CCARM 0230) to *Ampicillin* is shown in Fig. 5c. The MIC obtained via agar dilution method was eight μg/ml (Fig. 5d, Supplementary Movies 7, 8, and 9). The reverse peaks in the graphs for antibiotic concentration 0 and two μg/ml illustrates colony formations, while the eight μg/ml *Ampicillin* result shows the susceptible response of the bacteria. The sample of 4 μg/ml of antibiotic concentration, however, has a late peak at around *t* = 17.5 h with a large decrease of the correlation value, and this result still indicates the growth of bacteria according to the criterion. The different shape in the case of 4 μg/ml is analogous to a sample of lower initial concentration, which is described in Fig. 4b, and the relation between experiments of a high antibiotic concentration and a lower initial concentration can be explained by the optical images of the antimicrobial samples shown in Fig. 5d (Supplementary Movie 8 for detail). As can be seen, with *Ampicillin* of concentration of 4 μg/ml, only a small number of *E*. *coli* survives and develops into colonies, which is analogous to the lower initial concentration. This observation describes why the *C*(τ=7min) graph for *Ampicillin* 4 μg/ml is similar to the case of lower initial concentration of bacteria.

The results from antimicrobial susceptibility tests demonstrate a short time AST is available with the proposed speckle-based method. Typically, ASTs with agar dilution method take 16-20 h (2). This is mainly due to the time for colony development into a visible size and density. As investigated in Fig. 2, colonies are visible only after the exponential phase or reaching to the stationary phase. In the case of *E*. *coli* and *Ampicillin* in Fig. 5c, the growth of the well-growing samples (*Ampicillin* 0 and 2 μg/ml) almost finished before 14 hours, but the highly inhibited sample, *Ampicillin* concentration 4 μg/ml, slowly grows with the peak around at *t* = 18 h. This explains why the time 16-20 h is reasonable for the agar dilution method, because an early judgment before 14 hours could lead to an inaccurate MIC value of 2 μg/ml. However, with the proposed speckle-based method, the movement can be examined already at *t* = 12 h for the *Ampicillin* 4 μg/ml sample. To note, when observed with a microscope, a slight growth of bacteria seems to be observed even in higher *Ampicillin* concentration than MIC for initial a few hours (Supplementary Movie 9). This phenomenon can be explained as the result of a filamentary or swelling formation of bacteria in the presence of antibiotics (22). Fortunately, as can be seen in Fig. 5a, these unwanted dynamics did not cause a noticeable decrease in the graph. This indicates the sensitivity of the present optical setup conveniently selects colonial growth movements from other minor movements.

Here in the AST experiments, using the suggested criterion, the decreasing trend was identified before 4 hours in case of *S*. *aureus* to *Ampicillin*, and the slow bacterial growths of *P*. *aeruginosa* and *E*. *coli* with antibiotic concentration close to MIC were identified before 12 hours. Thus, applying the proposed method, the time for the susceptibility test can be reduced by 4-8 hours for *E*. *coli* + *Ampicillin* or *P*.*aeruginosa + Norfloxacin*, and by 12-16 hours for *S*. *aureus* compared to the agar dilution method, which takes 16-20 h, without compromising the accuracy. Of course, the visible development of colonies for some species may occur in a shorter time than 16-20 h, but the standard methods should take conservative criteria in order to prevent the misjudgment due to the slow growth of test samples. The present method provided fast and robust evaluation also for both the slow-growing, and even for the fast-growing samples, the decrease in correlation value occurred much before the visible colonies appear around the end of the peak time. In addition, the detailed antibiotic response of bacteria was also diagnosed as well as MIC, providing a deeper comprehension for the interaction between the bacteria and the microbial agents.

## Discussion and Conclusion

A new optical method for real-time bacterial growth detection based on speckle analysis has been proposed and experimentally demonstrated. We adopted a simple optical setup with two diffusers exploiting multiple light scattering, and investigated the samples by analyzing the temporal change of speckle fields. The proposed method was tested under various conditions of different bacterial species and concentrations, and the progress of colonial growth is well revealed providing a detailed history. It is for the first time that the precise characteristics of bacterial growth are identified on a solid surface, since, to the best of our knowledge, the only way to obtain an experimental growth curve is to measure optical densities of bacterial solutions in liquid form. Furthermore, the growth characteristics acquired via the present technique were distinguished by species implying the possibilities for species classification. For known species with calibration, we suggest that it is also possible to count the number of colonies from the sample of high-density solutions without dilution.

As the application of the present method, we demonstrated fast ASTs were possible by obviating the tedious time for visual development of colonies. This reduced the time required for the agar dilution method without compromising any part of the standard procedure or requiring additional treatment. Moreover, the simple configuration without lens system allows the freedom from focusing issues thus curtailing additional labor. Although the technique requires mechanical stability, we anticipate the simplicity and cost-efficiency of the method would offer a powerful alternative for current fast-AST tools.

## Methods

### Bacterial species and culture.

Bacterial species are chosen for each experiment. The experiment by species in Fig. 4a used three bacterial species *Escherichia coli* (*E*. *coli*, ATCC 8739, Korean Collection for Type Cultures (KCTC), Republic of Korea), *Pseudomonas aeruginosa* (*P*. *aeruginosa*, ATCC 9027, Korean Collection for Type Cultures, Republic of Korea) and *Staphylococcus aureus (S*. *aureus*, CCARM 3A168, Culture Collection of Antimicrobial Resistant Microbes, Republic of Korea). The same *E*. *coli* was used for the experiments in Figs. 2 and 3 ant the experiment by bacterial concentrations (Fig. 4b). For the antimicrobial susceptibility tests, three bacterial species but different strains, *Escherichia coli* (*E*. *coli*, CCARM 0230, Culture Collection of Antimicrobial Resistant Microbes, Republic of Korea), *Pseudomonas aeruginosa* (*P*. *aeruginosa*, CCARM 0219, Culture Collection of Antimicrobial Resistant Microbes, Republic of Korea), and *Staphylococcus aureus (S*. *aureus*, ATCC 29213) were used. All strains were purchased from the corresponding suppliers, except for *S*. *aureus* ATCC 29213 provided from *Samsung Medical Center*, Republic of Korea. The culture media for each species were adopted as listed on the species specification; *E*. *coli* and *P*. *aeruginosa* were cultured with Nutrient broth (ingredients purchased from BD Biosciences, USA), *S*. *aureus* CCARM 3A168 with Luria-Bertani broth (ingredients purchased from BD Biosciences, USA), and *S*. *aureus* CCARM 0219 with Tryptic Soy broth (ingredients purchased from BD Biosciences, USA). The manufacturing procedures strictly followed the protocol from KCTC.

Bacterial species were cultured overnight before use, and they were diluted to the desired concentration. The concentrations were adjusted measuring optical density (OD) with a spectrophotometer (UV-1280, Shimadzu Inc., Japan) at λ = 600 nm. The viable count of adjusted bacterial solution was confirmed through dilution and incubation on agar plates (49). Bacterial solutions were initially adjusted to 0.1 OD (1×10^8^ CFU/ml), and then diluted if needed, except for higher concentration adjusted to 0.5 OD (5×10^8^ CFU/ml).

### Agar plate and sample preparations

The procedure for making agar plates followed the protocol from KCTC. Appropriate culture broth components and agar powder (7.5 g for total volume of 500 ml) were mixed with distilled water (and antimicrobial agents if needed) and autoclaved. The solution at 70°C was distributed to petri dishes (40 mm in diameter, TPP 93040, Sigma-Aldrich, Germany), 3 ml for each. The agar plates were cooled and dried in a clean bench for 30 minutes while the fan is off. The manufactured agar plates were stored in a refrigerator at 4°C. For sample preparation, agar plates were taken out of the refrigerator just before their usage. The agar plates were dried in a clean bench (fan operating) about 10 minutes while adjusting the bacterial solution concentration. Then 5 μl droplet of the bacterial solution, or non-bacterial solution for a blank sample, was placed at the center of the agar plate. The agar plate was gently swayed in order to spread the bacterial area. The final bacterial area was circular and approximately 2 cm in diameter. Then the agar plates were dried in the clean bench (fan operating) for exactly one hour with its cover opened. After dry, the agar plates were covered and carefully mounted on the optical setup. Firmly fixed on the setup, the agar plates were maintained at 35°C with a custom-built temperature controller to provide a proper condition for bacterial growth.

### Antimicrobial agar plates.

For antimicrobial susceptibility tests, *Ampicillin* (A0166, Sigma-Aldrich Inc., USA) and *Norfloxacin* (N9890, Sigma-Aldrich Inc., USA) were chosen as test antimicrobial agents. The antimicrobial stock solution of concentration 100 mg/ml was prepared to dilute with phosphate buffered saline (PBS) and stored in a refrigerator at −20°C. The solutions were restored to room temperature before used and diluted to 10 mg/ml using PBS. The desired amount of the stock solution was added to the autoclaved agar solution at 70°C; for example, 1 μl of stock solution to 10 ml of agar solution for agar plates of 1 μg/ml antimicrobial concentration. Using 40-mm-diameter Petri dishes, the agar solution with antibiotics was solidified in a fan-off clean bench for 30 minutes with the cover ajar.

### Optical setup.

In order to achieve the detection of the bacterial growth movement on an agar plate, the optical setup was designed as the schematic shown in Fig. 1c. For the main vertical structure, 1.5-inch damped posts (DP14A, Lumenera Inc., USA) were used to achieve mechanical stability. A laser beam (λ = 635 nm, FLEXPOINT Dot Laser Modules®, LASER COMPONENTS Inc., Germany) was reflected by a 45 degree mirror (MRA25-E02, Thorlabs Inc., USA) and illuminated the sample vertically, while two diffusers (DG10- 1500-A, Thorlabs Inc., USA) were placed before and after the sample, respectively. The sample was placed in a custom-built aluminum molding with a temperature controller. After the second diffuser, a linear polarizer (LPVIS100, Thorlabs Inc., USA) was added for better contrast of the interference. Finally, a CCD camera (Lt365R, Lumenera Inc., USA) was mounted vertically facing down toward the sample. The size of a pixel was 4.54×4.54 μm^2^, and the whole image was 100×100 pixels. In order to adjust the speckle grain size, which depends on the numerical aperture of the system, an iris (SM2D25, Thorlabs Inc., USA) was added between the sample and the second diffuser. The size of a speckle grain on images was approximately 3×3 pixels, which corresponds to 14×14 μm^2^ on the image plane.

## Data availability

All relevant data are available from the authors.

## Acknowledgment

The authors appreciate the technical support from Chaegyun Jung

